# Neuromesodermal Progenitors are a Conserved Source of Spinal Cord with Divergent Growth Dynamics

**DOI:** 10.1101/304543

**Authors:** Andrea Attardi, Timothy Fulton, Maria Florescu, Gopi Shah, Leila Muresan, Jan Huisken, Alexander van Oudenaarden, Benjamin Steventon

## Abstract

During gastrulation, embryonic cells become specified into distinct germ layers. In mouse, this continues throughout somitogenesis from a population of bipotent stem cells called neuromesodermal progenitors (NMps). However, the degree self-renewal is associated with NMps in the fast-developing zebrafish embryo is unclear. With a genetic clone tracing method, we labelled early embryonic progenitors and find a strong clonal similarity between spinal cord and mesoderm tissues. We then followed individual cell lineages by light-sheet imaging and reveal a common neuromesodermal lineage contribution to a subset of spinal cord tissue across the anterior-posterior body axis. An initial population subdivides at mid gastrula stages and is directly allocated to neural and mesodermal compartments during gastrulation. A second population in the tailbud undergoes delayed allocation to contribute to the neural and mesodermal compartment only at late somitogenesis. We suggest that NMps undergo vastly different rates of differentiation and growth in a species-specific manner.

## Introduction

In amniotes, a bipotent population of neuromesodermal progenitors continually allocates cells to the posterior pre-somitic mesoderm (PSM) and spinal cord (Tzouanacou et al., 2009; Brown and Storey, 2000; Selleck and Stern, 1991). While no specific molecular marker has been identified for this population, they have been shown to express a combination of both the early neural and mesodermal markers Sox2 and Brachyury (Wymeersch et al., 2016), and have been named ‘neuromesodermal progenitors’ (NMps; Henrique et al., 2015). The observation of a NMp population is important as it suggest that germ layer specification continues throughout somitogenesis stages and is not restricted to primary gastrulation. Understanding when NMps allocate cells to spinal cord and paraxial mesoderm is an essential first step to explore the underlying molecular processes that determine the timing of germ layer allocation during vertebrate embryonic development.

The continuous allocation of cells from an NMp pool fits well with the cellular basis of axial elongation in mouse embryos, in which primary gastrulation generates only head structures and the rest of the body axis is generated by posterior growth (Steventon and Martinez Arias, 2017). However, externally developing embryos such as the zebrafish elongate their body axis in the absence of posterior volumetric growth, with elongation being a consequence of convergence and extension of mesodermal progenitors and volumetric growth of tissue within the already segmented portion of the body axis (Steventon et al., 2016). Despite these differences, genetic studies have revealed a marked conservation in the signal and gene regulatory networks that act to drive posterior body elongation across bilaterians (Martin and Kimelman, 2009). Furthermore, experiments in the zebrafish embryo have confirmed the presence of Sox2/Brachyury positive cells in the tailbud, and transplantation of single cells into the marginal zone can be directed to either neural or mesodermal cell fates depending on the level of canonical Wnt signalling (Martin and Kimelman, 2012). An additional progenitor pool exists within the tailbud that generates cells of the notochord and floorplate in a Wnt and Notch dependent manner (Row et al., 2016). While these studies demonstrate that a neuromesodermal competent population exists within the zebrafish tailbud, lineage analysis using intracellular injection of high molecular weight fluorescent dextran in zebrafish (Kanki and Ho, 1997) argues against a stem-cell like population that is homologous to the mouse NMp pool (Tzouanacou et al., 2009). These seemingly conflicting results can be resolved by a complete lineage analysis that determines the timing of neural and mesodermal lineage restriction in zebrafish, and answer in a clear manner whether these progenitors arise from a stem cell pool as they do in mouse embryos.

Here we use a genetic clone tracing method to address whether zebrafish NMps are a conserved source of spinal cord tissue, and the degree to which they populate neural and mesodermal structures during normal development. We find a closer clonal relationship between spinal cord and muscle as compared to spinal cord and anterior neural regions, which can be explained by a model of NM lineage decision at the basis of spinal cord generation in zebrafish. Tracing this lineage restriction with the combined use of photolabelling and the single cell tracking of lineages from an *in toto* light sheet imaging dataset demonstrate that this restriction occurs during an early and direct segregation event with little or none amplification of the cellular pool. We observe a second population of NMps that remains resident in the tailbud and contributes to the caudal-most portion of the tail, that matches a previously described tailbud NMp population (Martin and Kimelman, 2012). Taken together with recent studies, this suggests that an NMp population is a conserved source of spinal cord and paraxial mesoderm, but with large differences in their potential for self-renewal in vivo.

## Results

### Spinal cord scars are closer associated to mesodermal than to anterior neural derivatives

To determine whether a common NMp lineage contributes to the zebrafish body axis at a whole organism level, we used a novel CRISPR/Cas9 based genetic clone tracing method called ScarTrace that allows for the reconstruction of clonal relationships in a retrospective manner (Alemany et al., 2018; Junker et al., 2016). When Cas9 RNA or protein is injected together with a guide RNA (gRNA) targeting a tandem array of histone-GFP transgenes in a zygote, a series of insertions and deletions of different lengths at different positions (scars) are introduced subsequently to DNA double strand break. These scars are inherited to all daughters of each labelled progenitor cell.

If a NMp population is at the root of the spinal cord lineage in zebrafish, it would be expected that descendants of the paraxial mesoderm tissues would have similar scars to those within the spinal cord. Alternatively, if spinal cord were generated from an ectodermal territory that is segregated from the mesoderm prior to anterior/posterior neuronal lineage segregation as predicted by the activation/transformation model (Niehrs, 2001; Nieuwkoop and Nigtevecht 1954), it would be expected that it would share a common set of scars with brain regions. To distinguish between these two possibilities, we injected either Cas9 RNA (fish R1 to R3) or Cas9 protein (fish P1 to P3) together with guide RNAs against GFP at the 1-cell stage into embryos transgenic for 8 copies of H2A-GFP (Figure 1A). Cas9 RNA injection has been shown to generate scars up until 10 hpf and therefore would label any neural and mesodermal progenitors throughout gastrulation, while Cas9 protein scarring ends at around 3hpf (Alemany et al., 2018). At larval stages, multiple regions of the body were isolated mechanically and sequenced to determine their scar composition (Materials & Methods). To determine the scar-based distance between organs we used hierarchical clustering (Figure 1B, Figure S1A,B). Importantly, a strong relationship was observed between spinal cord and muscle tissues at all anterior-posterior (labelled front, mid and tail) regions of the body axis with Cas9 mRNA injections, suggesting that at least by 10 hpf, a common neuromesodermal population underlies spinal cord development at all axial levels. For Cas9 protein injected fish we observe a closer association of spinal cord tissue across the anterior to posterior axis than to muscle tissue (Figure 1 –Figure S1B). However, due to the absence of muscle location in P1-P3 it can either be that we label spinal cord progenitors before mixing with muscle or that muscle cells only in close proximity to the spinal cord have a shared progenitor history. This observation suggests two models for the segregation of spinal cord and paraxial mesoderm fates during zebrafish embryogenesis. The first model, continuous allocation, follows the interpretation of retrospective lineage analysis in the mouse (Figure 1C) and assumes that both spinal cord and paraxial mesoderm cells are continually produced from a posteriorly localised neuromesodermal stem cell pool as in mouse. The alternative is the early segregation model, according to which neural cell lineages diverge during early gastrulation stages concomitant with the early specification of mesoderm tissue (Figure 1D). To determine the relative contribution of these two segregation models of zebrafish NMps, we turned to imaging-based lineage tracing methods.

**Figure 1.**
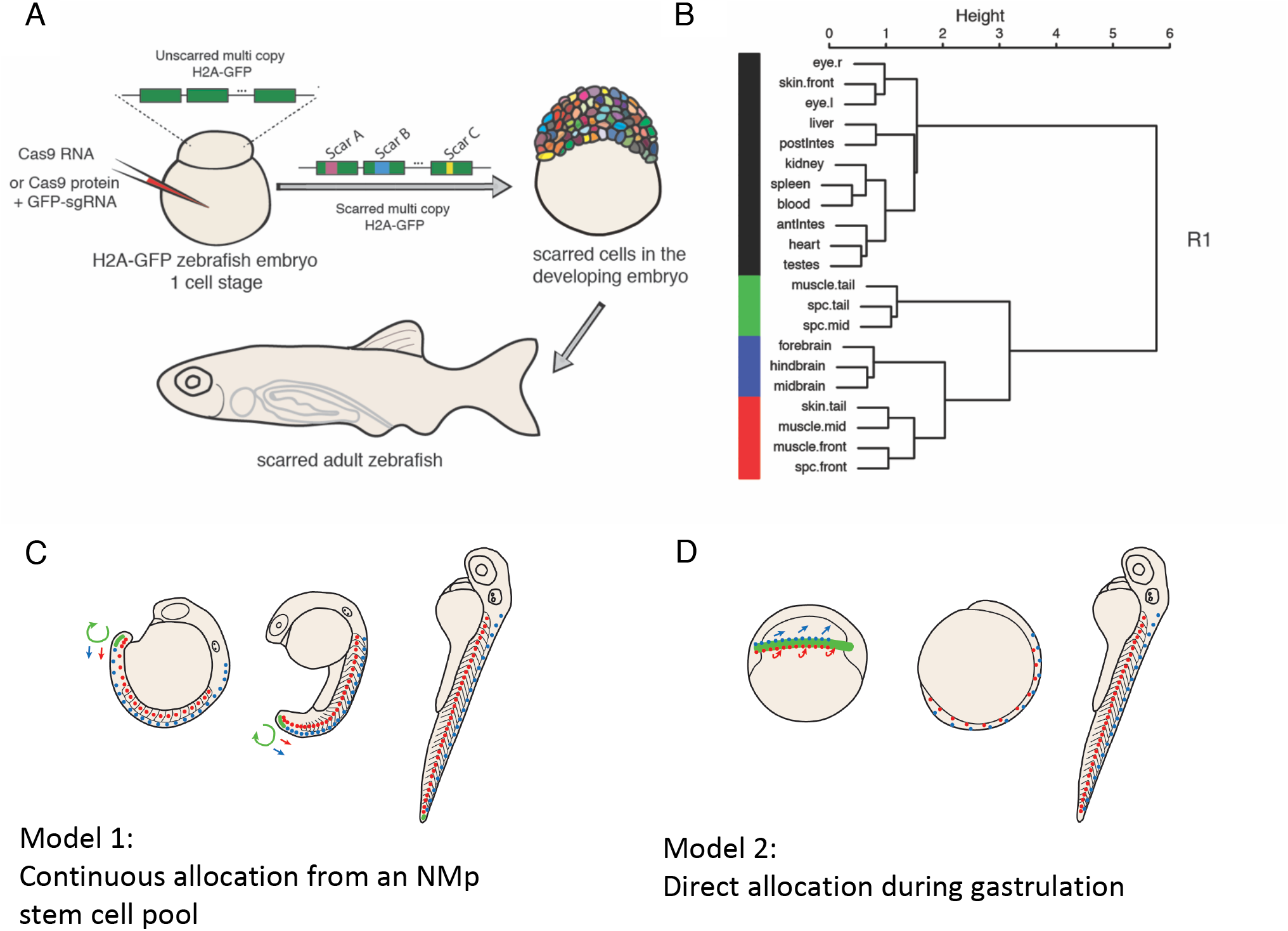
Spinal cord clones are closely associated to mesodermal and not anterior neural derivatives. A) Experimental workflow for Scartrace lineage tracing of the zebrafish larvae. B) Dendrogram showing the relationships between dissected larval body structures in R1. The height of the dendrogram shows the hierarchical clustering of Canberra distances between the spearman correlation of organ scar fractions. The colours show the four most distant branches. Note the close association of spinal cord and mesodermal cells across the anterior to posterior axis, that is distinct from brain regions, antIntes = anterior intestine, postlntes = posterior intestine, l. = left, r. = right, Spc. = Spinal cord C) The continuous allocation model is based on lineage analysis in mouse embryos and supposes that a group of self-renewing neuromesodermal stem cells continually generate spinal cord and paraxial mesoderm throughout axial elongation. D) The direct allocation model proposes that an early segregation of neural and mesodermal lineages occurs during gastrulation, the derivatives of which then expand rapidly by convergence and extension.

### Tissue level segregation of spinal cord and mesoderm clones occurs by 50% epiboly

To obtain an estimate of when spinal cord and paraxial mesoderm spatially segregate during zebrafish gastrulation, we performed a series of fate mapping experiments using photolabelling. Embryos were injected at the one-cell stage with mRNA encoding for the photoconvertible protein, Kikume, targeted to the nucleus (nls-Kikume). Upon exposure to UV light at early gastrula stages, regions of around 50-100 cells spanning the marginal zone were labelled, enabling the direct visualization of mesoderm and ectoderm segregation by confocal microscopy. While labels in the prospective anterior-dorsal region of the 30% epiboly embryo contributed to large portions of the brain and notochord (Figure 2A), these labels contributed little to the spinal cord and paraxial mesoderm territories (Figure 2D), supporting the distinct lineage for brain regions found by Scartrace (Figure 1A,B). The co-labelling of anterior neural and notochord fated cells reflects the fact that the prospective shield region at 30% epiboly partially overlaps with the labelled ectodermal domain. In contrast, labels along the medial portions of the marginal zone generate significant contributions to both spinal cord and paraxial mesoderm compartments (Figure 2B,E). Labels in an equivalent domain at 50% epiboly resulted in significantly less mesodermal contributions, in line with the continued invagination of mesodermal progenitors at these stages (Figure 2C,F). However, not even small (36 cells) labels could mark solely mesoderm fated cells, suggesting that a degree of mixing between ectodermal and mesodermal lineages persists until 50% epiboly (Figure 2G).

Following the 50% spinal cord/mesoderm fated clones by time-lapse microscopy reveals a rapid convergence and extension of spinal cord progenitors that lead to a widespread contribution across a large proportion of the anterior-posterior axis (Supplementary movies 1 and 2). Indeed, labelling of a single region in a medial position at 50% epiboly stage results in the labelling of the neural tissue from the base of the hindbrain to the tailbud at the 16-somite stage (Figure. 2E). Cells that remain ectodermal upon invagination of the mesoderm become displaced posteriorly by the continued convergence and extension of cells in the animal pole (Supplementary movie 3). Thus, it appears that a large proportion of the spinal cord is allocated during gastrulation stages, and that this arises from a domain close to or overlapping with a paraxial mesoderm domain. However, without single cell lineage tracing, it is not possible to conclude whether these cells arise from a common neuromesodermal progenitor pool.

**Figure 2.**
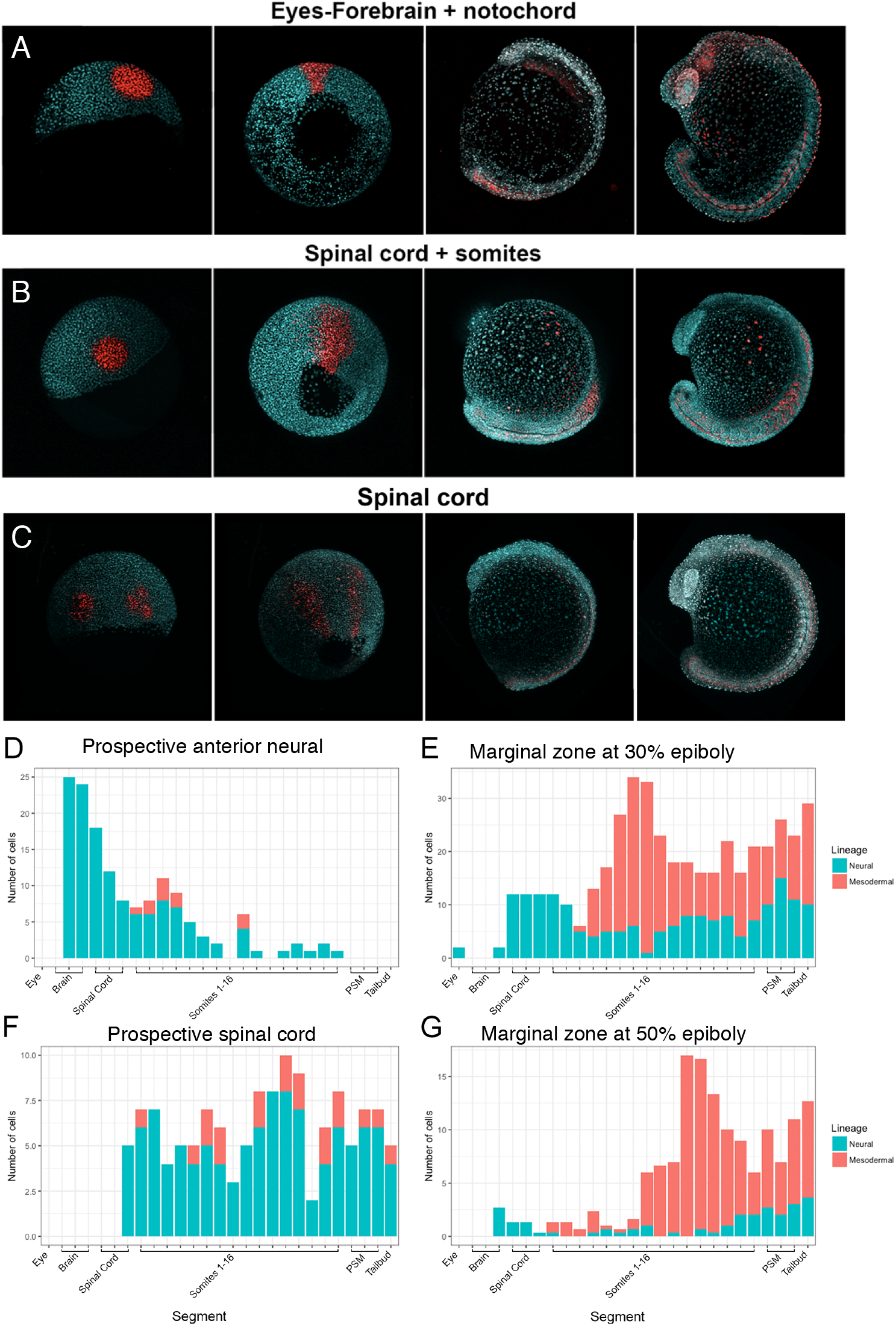
Tissue level segregation of spinal cord and mesoderm clones occurs by 50% epiboly. (A-C) Embryos were photolabelled in the prospective anterior neural (A), marginal zone (B) and prospective spinal cord (C) territories at 30% epiboly (left-most image) and followed until the 18 somite stage (right-most image) by time-lapse microscopy. (D-G) The contributions of labelled clones from individual examples are plotted against the anterior-posterior axis with the number of cells in each tissue compartment shown in red for the somitic mesoderm or blue for the neural tube. Note that a significant degree of overlap between spinal cord and mesoderm fated cells exists within the marginal zone at both 30% (E) and 50% (G) epiboly.

### Bipotential neuromesodermal cells segregate rapidly during mid to late gastrulation

To assess whether single cells contribute to both spinal cord and mesoderm we made use of an existing light sheet dataset in which the onset of mesoderm specification can be observed with the use of a live reporter for *mezzo* (Shah et al., 2017). In this dataset, germ layer segregation can be assessed live by detecting the increase in mezzo:eGFP fluorescence levels in the nuclei of mesendodermally specified cells (Figure 3A). Segmentation and automated tracking of all nuclei within the gastrulating embryo, a custom Matlab script was used to isolate the entire tracks of cells in a user-defined region of the embryo at a chosen time point, allowing us to perform labelling experiments *in silico*. Automated cell tracking performed using TGMM (Amat et al., 2014) was validated by manually inspecting each track using the Fiji plugin Mamut (Wolff et al., 2017).

To characterize the germ layer allocation of individual cells within the medial marginal zone at 30% epiboly, we focused on a region comparable to the clone of cells fated towards both spinal cord and paraxial mesoderm at 30% epiboly (Figure 2B,E). We hypothesized that the more animal part of this region could consist of bipotent progenitors. In fact, separation of the tracks of marginal cells from the rest of cells present in the medial region shows that overall track lengths of marginal cells are shorter than those of more animal cells, all ending before shield stage (Figure 3B). Shorter track length of these cells is a consequence of their rapid mesodermal segregation though marginal involution (Supplementary Movie 4). We then focused on the more animal cells of the clone, whose continued up to the end of epiboly, and allocated their fate as being either spinal cord or mesoderm (n=100) (Figure 3B). Cells which retained ectodermal lineage at track termination contributed to the spinal cord (Figure 3, Figure S2) and are shown with a blue bounded box in Figure 3C. Those that increased in the expression of mezzo:eGFP were determined as being of mesoderm fate (Supplementary movie 4) and have a red bounded box (Figure 3C). Within this region of the marginal zone, 68 trajectories retained ectodermal specification in all their daughter cells at the termination of their tracks with the remaining 23 undergoing specification to mesoderm. Importantly, seven tracks contributed to both the mesodermal compartment and the spinal cord. Two tracks were excluded due to gross inaccuracies in tracking. 75% of tracks did not undergo an identifiable cell division over the tracking period. Based on this, we conclude that a small population of bipotent neuromesodermal progenitors exist close to the marginal zone of the zebrafish embryo, but rapidly segregate between the 70-90% epiboly stage and are mixed with largely monopotent progenitors.

**Figure 3.**
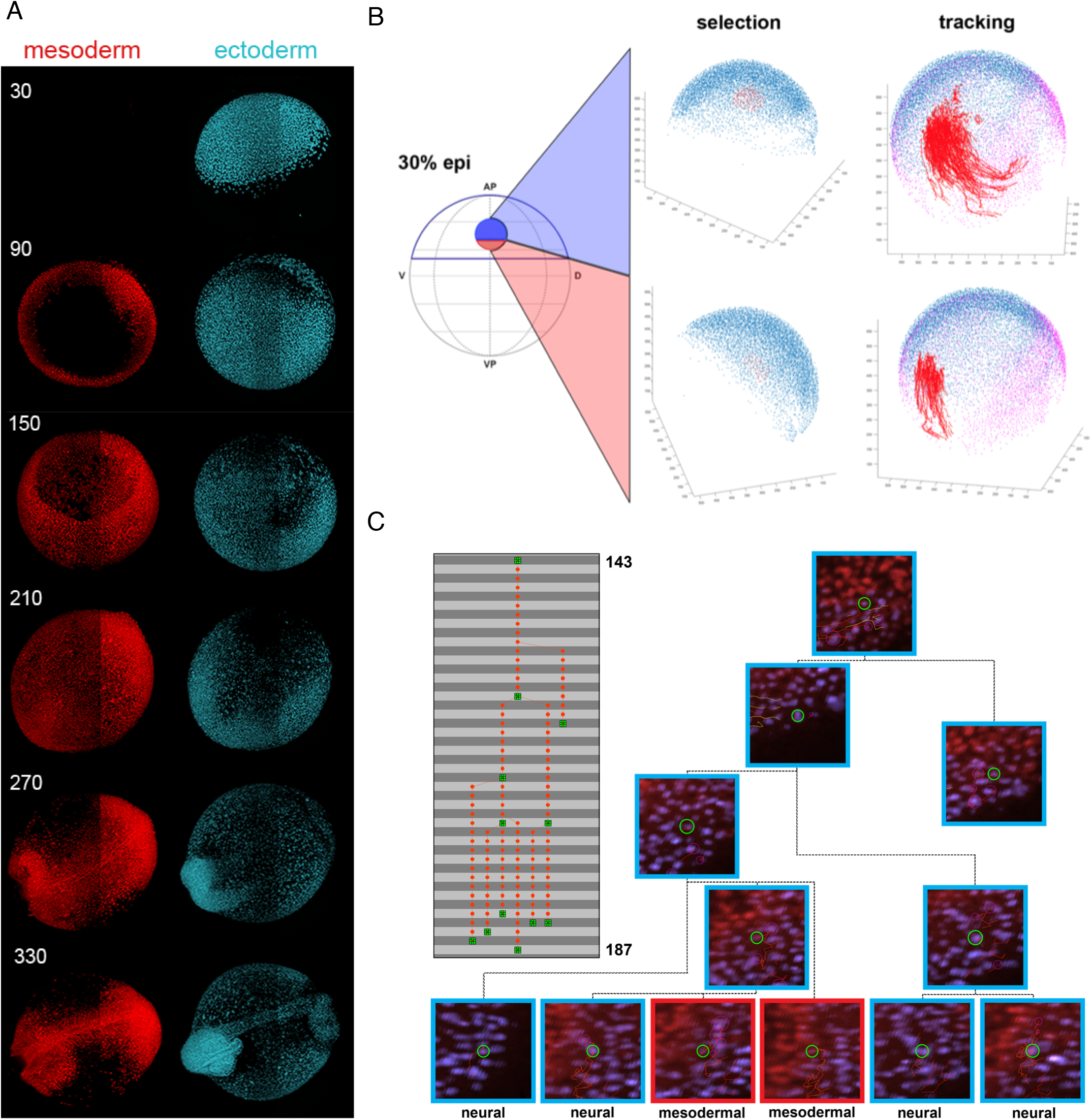
Bipotential neuromesodermal cells segregate rapidly during mid to late gastrulation. A) Stills of a multi-view light sheet movie (Shah et al, 2017), show the progressive onset of mezzo:eGFP expression (channel depicted in red) as cells become committed to the mesendodermal lineage. Numbers on the left refer to the time point of acquisition. B) A custom MATLAB-based tool was developed to allow for the extraction of tracking data from the marginal zone at 30% epiboly. Tracks were selected separately (n=100) in marginal and animal subgroups and visualised on top of 75% epiboly stage nuclei as a reference (purple). C) Tracks containing cells that give rise to both spinal cord and mesodermal derivatives (n=7). A 45 timepoint segment of a representative lineage of cells in the upper marginal zone is shown in the grey box. Cells highlighted in green are shown in the tree on the left, with dashed lines indicating indirect relationships.

Blue and red bounding boxes indicate the germ layer allocation of individual cell as they progress along their tracks.

### Tailbud neuromesodermal clones are restricted to the posterior most aspect of the body axis

Upon completion of gastrulation the tailbud forms via a continued convergence and extension of cells from the anterior ectoderm and a concomitant subduction of posterior cells to form the paraxial mesoderm (Kanki and Ho, 1997). The posterior-most subset of the labelled gastrula-stage NMp population becomes located within the dorsal tailbud upon the completion of primary gastrulation (Supplementary movie 3), suggesting that there may be a spatial continuity with the Sox2/Brachyury positive population in this region (Martin and Kimelman, 2012). We used conventional fate mapping to assess the contribution of this population. Using nls-Kik injected embryos, multiple regions of the tailbud were photolabelled at the 6-somite stage and then reimaged at the end of somitogenesis; an intermediate image was taken at the 22-somite stage. A progenitor region which give rise to both neural and mesodermal tissue are identified dorsal to the posterior end of the notochord (Figure 4A). In tracking these cells, it was shown that between 6 and 22 somite stages the progenitor cells track the dorsoposterior end of the notochord and showed no contribution to the extending axis. The population was also observed to undergo a dorsal-to-ventral rotational movement along the posterior wall, aligning with the caudal neural hinge. The NMp clones were observed only contributing to the final seven somites (25^th^ to 32^nd^) and nine neural segments (23^rd^ to 32^nd^).

Directly adjacent anteriorly to the NMps, a progenitor pool with only neural fate was identified. This population contributed to earlier forming spinal cord and no mesodermal tissue. Labelled neural clones were shown to contribute to neural segments between the 14^th^ and 27^th^ somites. No mesoderm was labelled in these clones. Directly adjacent posteriorly to the neural mesoderm progenitors, a third population of progenitors was identified which only contributed to mesodermal tissues, in particular the earlier forming somites between 19^th^ and 31^st^.

To check whether our labels are covering the Sox2^+^Ntla^+^ domain previously described (Martin and Kimelman, 2012), we compared the labels to HCR stains for these genes (Choi et al., 2014; Figure 4F). By segmenting each expression domain, the region of overlap can be visible (Figure 4G; Supplementary movie 5). The *ntla* expression domain was used to mask all *sox2* expression outside of this region of interest, thereby allowing only regions with co-expression to be observed in 3D (Figure 4H; Supplementary movie 6). This region is overlapping with the example NM labelled population shown in (Figure 4A,I).

In labelling of the early progenitor populations and identifying their final fates at the end of somitogenesis (Figure 4 A-E), the number of cellular divisions was calculated (Figure 5). Neural and mesoderm specific progenitor populations were shown to both have a significantly (p<0.001, n=7) higher rate of proliferation than the neuromesodermal clones which were shown to replicate infrequently. The monopotent clones (MPs and NPs) underwent a much higher frequency of cell divisions with NPs doubling in cell number between bud stage and the end of somitogenesis.

**Figure 4.**
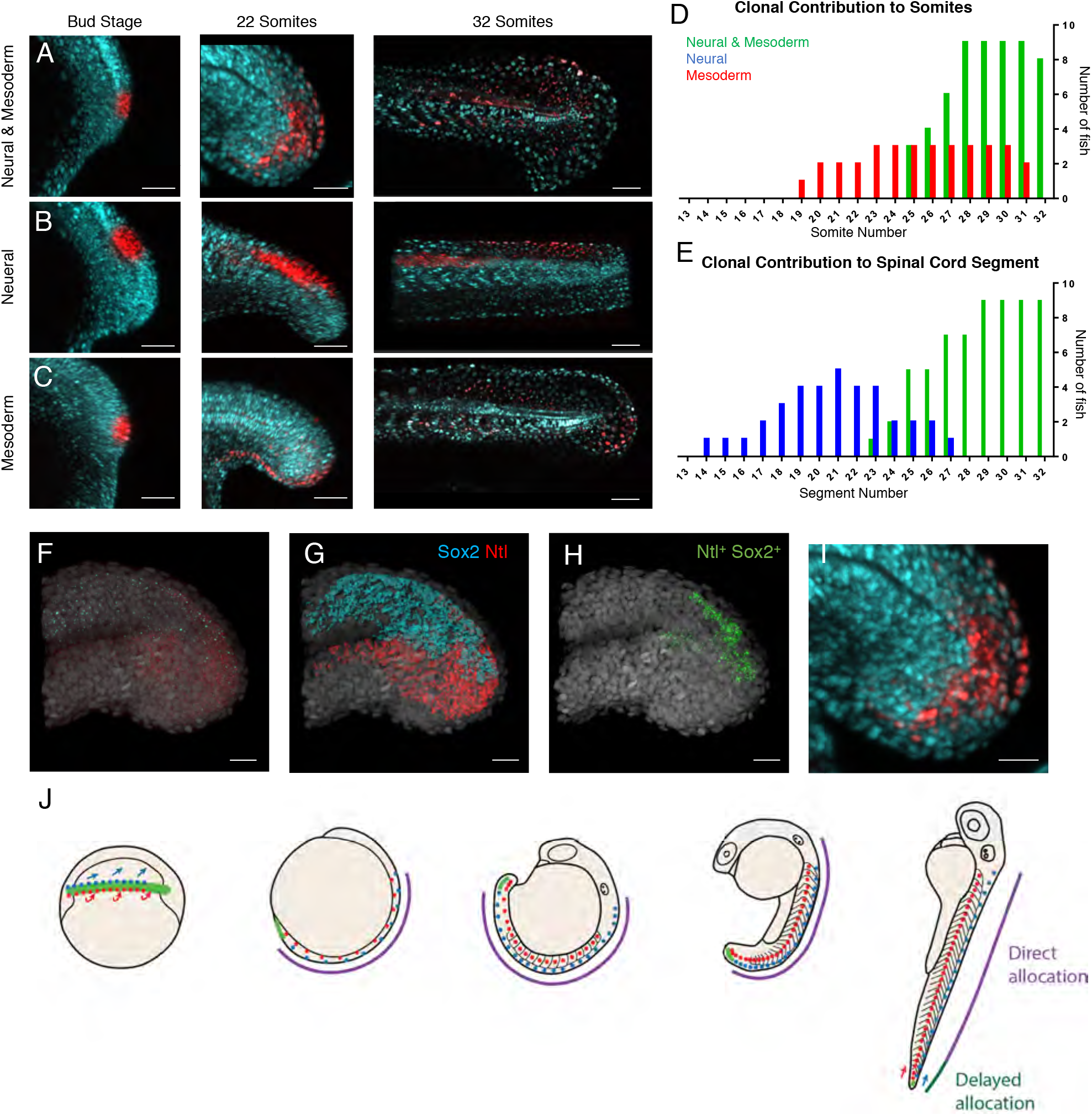
Tailbud neuromesodermal clones are restricted to the posterior most aspect of the body axis. (A-C) Regions of the tailbud where photolabeled at bud stage and followed until the completion of somitogenesis. Populations on the dorsolateral wall of the tailbud gave rise to both spinal cord and mesoderm derivatives (A), those more anterior generated only spinal cord (B) and those more ventral only to mesoderm (C). All labels had an additional contribution to non-neural ectoderm. The contribution of clones to the anterior-posterior axis are plotted separately for mesodermal (D) and neural (E). In red are labels with mesodermal contribution only (n=7), in blue are spinal cord specific clones (n=5) and in green are those that gave rise to both germ layers (n=9). (F-I) Hybridisation Chain Reaction (HCR) for *sox2* and *ntla* was used to locate double positive cells within 3D confocal datasets. (F) Original dataset with *sox2* in cyan and *ntla* in red, DAPI is in grey. (G) Surface segmentation of the HCR stain was performed to mask *sox2* expression within the *ntla* channel. (H) Masking reveals only those cells that are co-expressing both genes. (I) Magnified image of that shown in (A) to compare photolabel with co-expressing cells.

**Figure 5.**
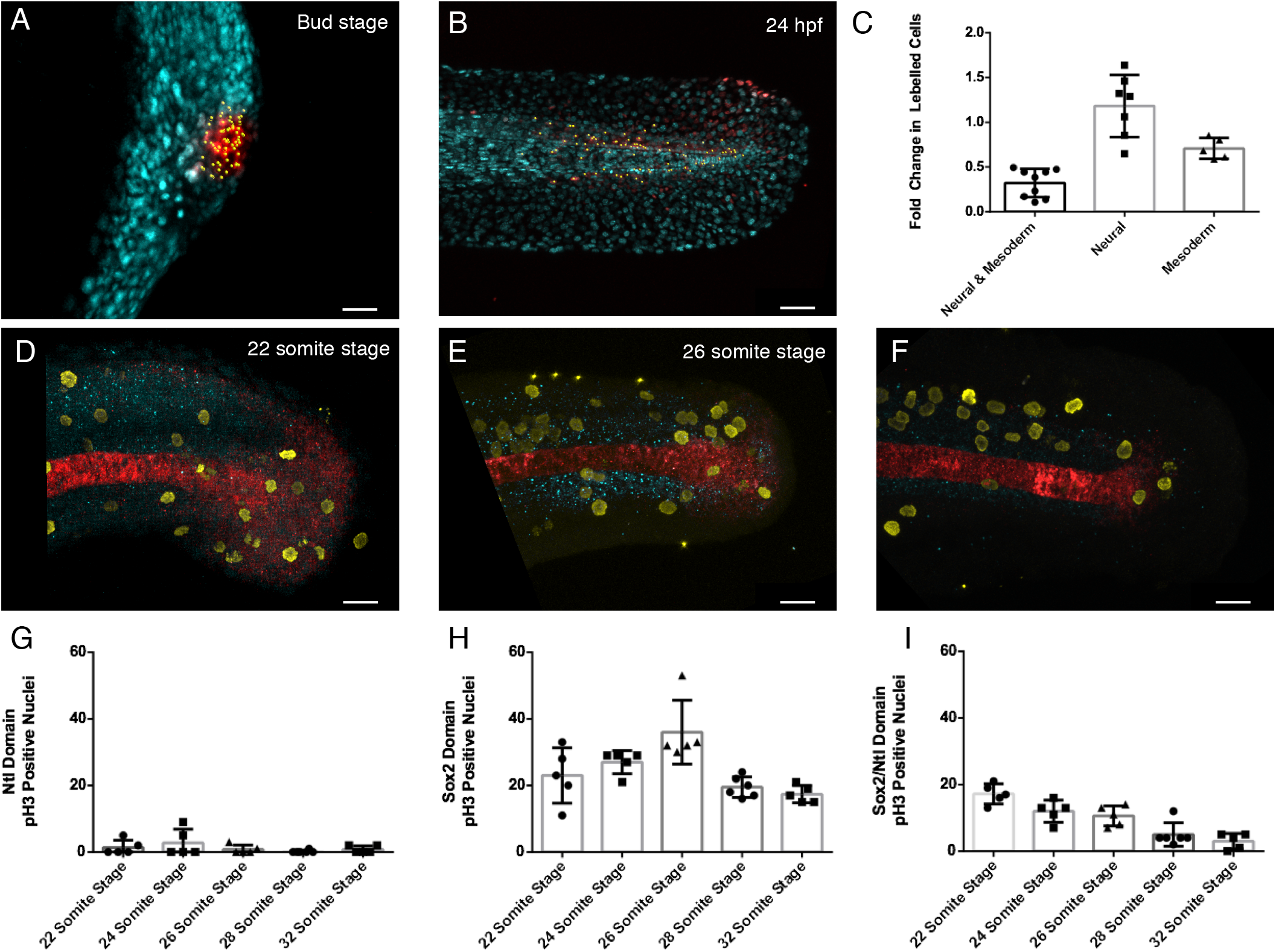
Quantification of cell division in tailbud NMps. Quantification of increase in number of cells photolabelled using nls-kikume from (A) bud stage through to (B) 24hpf. (C) Fold change increase in labelled clone number changes depending on labelled progenitor type with clones contributing only to neural tissue undergoing most clonal expansion and bipotent progenitors undergoing the least clonal expansion. (D-F) Replicating cells stained using phosphor-histone H3 (pH3) as a marker of mitotic cells (yellow) with bipotent NMps identified through coexpression of Sox2 (blue) and Ntl (red) at the (D) 22 somite stage (E) 26 somite stage and (F) 32 somite stage. The frequency of Ph3 positive nuclei in the (G) Ntl positive expression domain including notochord, (H) Sox2 positive expression domain and (I) Ntl and Sox2 positive NMp domain.

## Discussion

Taken together, these results show that a neuromesodermal progenitor pool is a source for at least a subset of spinal cord cells at all regions of the body axis. Critically however, this is not achieved by a bipotent stem cell population but rather by the rapid segregation of neural and mesodermal cell fates during gastrulation. Subsequently, a second population of neuromesodermal progenitors arise along the dorsal wall of the tailbud, co-expressing both *sox2* and *ntla* (Martin and Kimelman, 2012; this study). However, these are not continually added to the elongating body axis as previously supposed but are rather a largely quiescent population of cells that contribute to the last portion of the body axis. The low proliferation levels of these cells during somitogenesis are in line with a previous study (Bouldin et al., 2014). While it is possible that a proportion of cells are bipotent within this population, the low rates of division make it unlikely that their derivatives will have sufficient time prior to the completion of somitogenesis to generate a stem-cell mode of growth as has been observed in mouse embryos (Tzouanacou et al., 2009). Therefore, we surmise a composite model for spinal cord generation in zebrafish with spatially and temporally separate pools. The first is directly allocated during gastrulation and is a population of NMps mixed in with a large proportion on monopotent progenitors. The second matches with a previously described tailbud NMp population (Martin and Kimelman, 2012) and has a delayed allocation to only the final portion of the larval tail (Figure 4J).

Early studies of vertebrate axial elongation by D.E. Holmdahl in the mid-1920s, lead to the suggestion of a distinct ‘secondary body development’, that is distinguished from the early gastrulation process by the continuous generation of cells from a homogeneous blastemal (Holmdahl, 1925a,b,c; Handrigan). Walther Vogt saw the process differently and postulated that secondary body development proceeds as a continuation of gastrulation (Vogt 1926). This alternative view has been supported by the observation that the Xenopus tailbud contains germ layer restricted progenitors and tissue specific localisation of Bracyhury (Tucker and Slack 1995a,b; Gont et al., 1993). As NMps continue to generate multiple germ layer derivatives throughout secondary body development, one might argue that this supports a constrained version of Holmdahls’ blastemal hypothesis, as the tailbud contains cells that are not yet restricted in their germ layer potential. However, as NMps are also present through gastrulation stages, the fact that they continue to undergo a similar cell fate decision within the tailbud suggests that secondary body development is very much a continuation of gastrulation. Whether a similar gene regulatory network underpinning the specification of cells from both these early and late NMp populations remains an open question.

Despite a strong developmental constraint acting upon gastrulation in vertebrates (Abzhanov, 2013), there exists a large degree of morphological variation acting at these stages of development (Duboule, 1994). This variation is largely a consequence of the different strategies of maternal-embryo energetic trade-off that have been adopted during chordate evolution. In the context of mouse, viviparity has led to an internal mode of development within which embryos increase in volume to a large degree concomitantly with establishing their body plan (Steventon et al., 2016). This is in contrast to macrolecithal embryos such as fish, whose energy supplies are contained within an external yolk sac. Such transitions have a great impact on the morphology of gastrulae, that must adopt their shape according to the physical constraints of yolk size and extraembryonic structures. Furthermore, external modes of development provide a selective advantage for developmental strategies that favour a rapid development to swimming larval stages in order that they escape predators and find food. Zebrafish undergo a highly rapid mode of development, and develop their full complement of somites, prior to the development of a vascular system that is efficient at accessing nutritional supplies and allowing them to increase in mass. We propose that by shifting the allocation of spinal cord and mesodermal progenitors to early gastrulation stages, this has facilitated a rapid convergence and extension-based mode of axial elongation. Interestingly, the pool of NMps within the tailbud demonstrates a conservation of a tailbud progenitor pool that could allow for increased flexibility according to organism level heterochronies in the rates of growth. This hints at a novel evolutionary developmental mechanism that we term ‘growth-mode adaptability’. This proposes that specific cellular trajectories (such as the emergence of spinal cord from a neuromesodermal cell state) are conserved in evolution, yet highly adaptable in terms of their timing of cellular decision making and modes of growth.

## Supplementary Material

**Figure S1.**
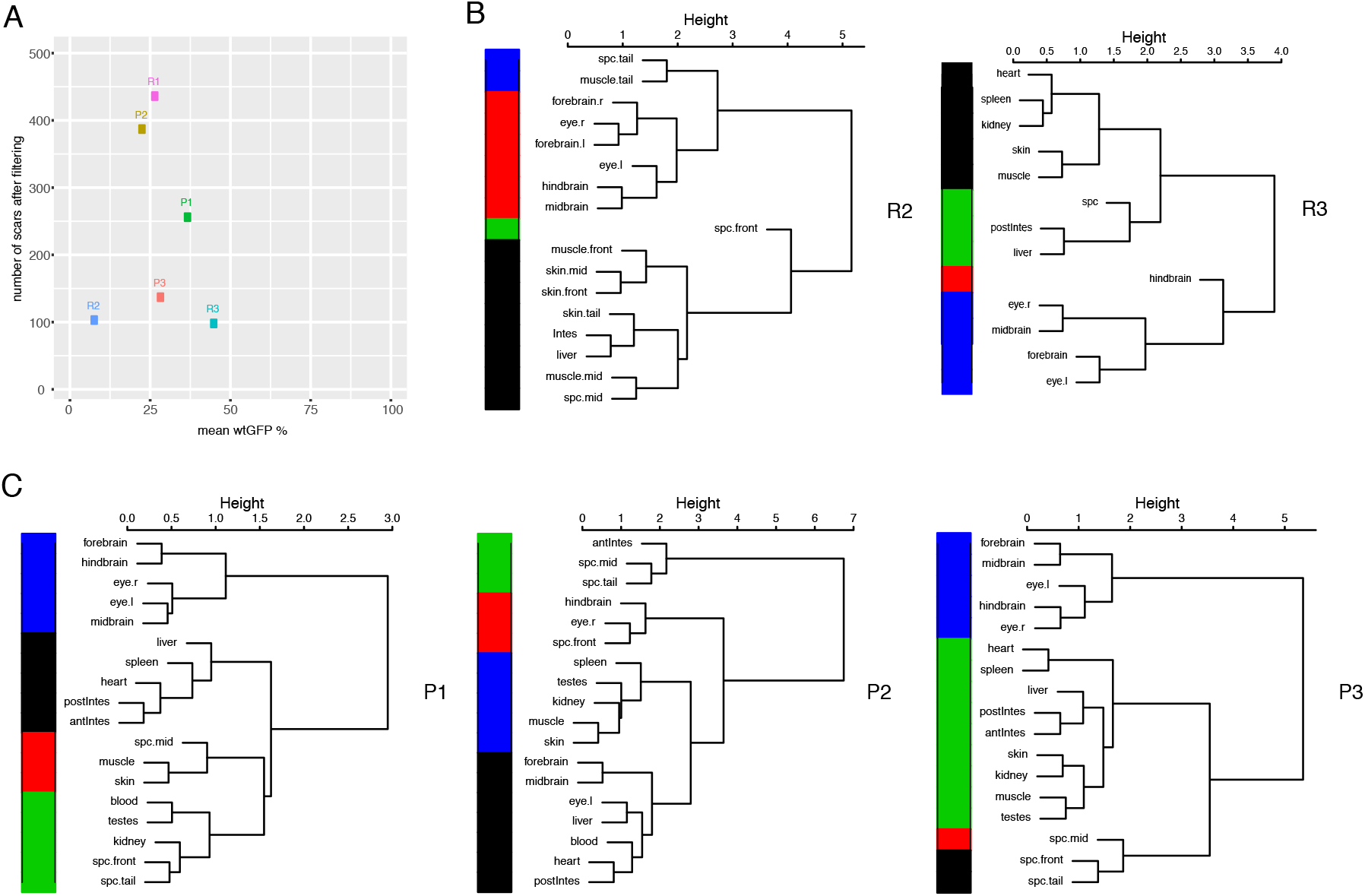
ScarTrace in Cas9 RNA or protein injected fish. (A) Replica R2 and R3 for Cas9 RNA injected fish. The height of the dendrogram shows the hierarchical clustering of canberra distances between the spearman correlation of organ scar fractions. The colours show the four most distant branches. Note the close association of spinal cord and mesodermal cells across the anterior to posterior axis, that is distinct from brain regions. antIntes = anterior intestine, postIntes = posterior intestine, l. = left, r. = right, Spc. = Spinal cord (B) Dendrograms as described in (A) for Cas9 protein injected fish P1 to P3. (C) Scarring efficiency shown as the amount of unique scars in all organs after filtering vs. the mean unscarred GFP percentage per fish. Only fish with less than 50% unscarred GFP (uGFP) were used. R refers to Cas9 RNA injected and P to Cas9 protein injected fish.

**Figure S2.**
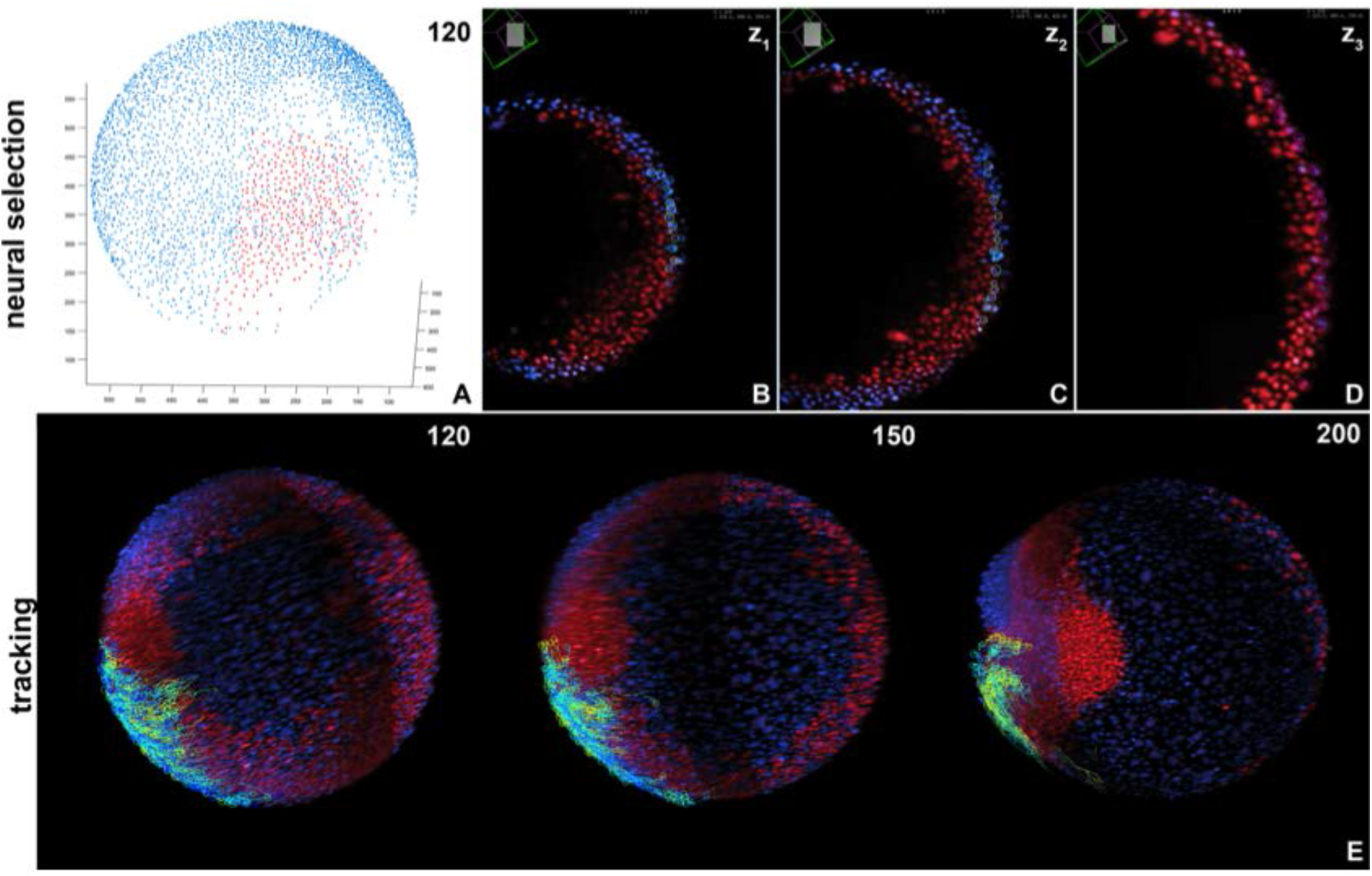
Spinal cord allocation of upper marginal cells. (A) Cells of the upper marginal label shown in Figure 3 are selected (red spots) and followed from timepoint 120. (B-D) Mamut Viewer visualisation of the selection at three different optical sections shows that cells lack *mezzo* expression and are still localised in the outer layer of the blastoderm. (E) Further tracking of these ectodermal cells shows their migration towards the embryonic anteroposterior axis and final arrangement in one of the two arcs of cells that converge to form the spinal cord. Images in E depict partial maximum intensity projections to exclude the head region at the indicated timepoints.

## Materials and Methods

### Animal lines and husbandry

This research has been regulated under the Animals (Scientific Procedures) Act 1986 Amendment Regulations 2012 following ethical review by the University of Cambridge Animal Welfare and Ethical Review Body (AWERB).

### Scartrace

Scarred zebrafish (injected with RNA Cas9 or protein Cas9 and gRNA against GFP) were generated as described in Junker et.al, 2016 and Alemany et al. 2018. Scarred zebrafish were anaesthetised, sacrificed and small parts of organs were dissected and collected individually. Genomic DNA was isolated from the organ parts using DNeasy Blood & Tissue Kits (Qiagen, Cat No./ID: 69506) and scars were amplified and sequenced using the ScarTrace bulk protocol as described in Junker et al., 2016. Scars were mapped as described in Alemany et.al, 2018. After mapping, to filter out potential sequencing errors, the scar percentage was computed in normalized histogram with 100 bins. Scars were only kept when their fraction was at least 10x higher than the minimum detected scar fraction. In case multiple organ parts of the same tissue were dissected the average scar fraction was computed. To filter out fish with inefficient scarring we determined the scarring efficiency as the number of scars after filtering and the mean unscarred GFP percentage across all organs per fish (Figure S1C). Only fish with less than 50% unscarred GFP and more than 100 scars were kept. Additionally, only unique scars (not occurring in other scarred fish) were used to compute the dendrograms.

### Preparation and mounting of zebrafish samples for photolabelling

Zebrafish wild-type embryos were injected at the one-cell stage with 200 pg of nuclear-targeted Kikume (kind gift of Ben Martin). Photolabelling was performed as described in (Steventon et al., 2016). For labelling, embryos were mounted in 2% methycellulose in E3 and low gelling point agarose in E3, described in (Hirsinger and Steventon, 2016), for live imaging of post gastrulation embryos.

### Hybridisation chain reaction and immunohistochemistry

Five antisense DNA probes were designed against the full length zebrafish *sox2* and *ntla* mRNA sequence as described in Choi et al. (2014). Embryos incubated in either embryo medium were fixed 4% PFA at 4°C and then stained according to Choi et al. (2014). HCR was combined with immunohistochemistry for phospho-histone H3 for identification of mitotic cells at the point of fixation. Embryos underwent HCR followed by blocking in 2% Roche Blocking reagent, 5% donkey serum in MAB. A mouse anti-pH3 antibody (abcam, ab14955) was incubated overnight at 4 degrees followed by a secondary anti-mouse conjugated to an Alexa 488 fluorophore (ThermoFischer Scientific, A32723). Imaging was performed on a Zeiss LSM700 confocal with identical imaging parameters: z-step 0.5683μm; 1024×1024 resolution; 63X oil objective; 35% laser power at 488 nm; Gain 661; Digitial offset −2; Pixel dwell time 3.12μs.

### Matlab script for trajectory selection

The custom written Matlab script imports trajectories saved in TGMM software format (Amat et al., 2013) and visualizes the position of the detected cells at two user defined points in time. Through GUI the user selects the cells of interest and subsequently the trajectories of the respective cells are plotted in red. An (optional) post-processing step eliminates the cells on the opposite side of the embryo from the user’s viewpoint, in case they were inadvertently selected. The selected trajectories can be saved in the TGMM software format or as a Matlab “.mat” file.

### *In silico* fate mapping

The light sheet dataset used for the lineage analysis, as well as the TGMM automated tracking, is described in Shah et al., 2017. Trajectories selected using our Matlab script at timepoint 30 (corresponding to 30% epiboly) were exported as TGMM output xml file. TGMM data has been imported in MaMuT (Wolff et al., 2017) for visualization and validation of tracks. Lineages were followed using the Track Scheme module and inspected by overlaying the tracking data on top of the original images in Mamut viewer. Germ layer segregation of cells along tracks was determined based on the expression of the *mezzo:eGFP* mesodermal reporter.

## Acknowledgments

We would like to thank Alfonso Martinez-Arias, Estelle Hirsinger, Octavian Voiculescu and all members of the Steventon lab for comments on the manuscript. We also kindly thank A. Alemany for input on the ScarTrace analysis and the manuscript. B.S. and T.F. are supported by a Henry Dale Fellowship jointly funded by the Wellcome Trust and the Royal Society (109408/Z/15/Z). A.A. was supported by the “Erasmus+ for Traineeships” scheme of the European Union. Work in the lab of J.H. was supported by the Max Planck Society and European Research Council (CoG SmartMic, 647885). The work of M.F. and A.V.O work was supported by a European Research Council Advanced grant (ERC-AdG 742225-IntScOmics), Nederlandse Organisatie voor Wetenschappelijk Onderzoek (NWO) TOP award (NWO-CW 714.016.001), and the Foundation for Fundamental Research on Matter, financially supported by NWO (FOM-14NOISE01). This work is part of the Oncode Institute which is partly financed by the Dutch Cancer Society.

## References

Abzhanov, A. (2013). von Baer’s law for the ages: lost and found principles of developmental evolution. Trends Genet. 29, 712–722.

Alemany, A., Florescu, M., Baron, C. S., Peterson-Maduro, J. and van Oudenaarden, A. (2018). Whole-organism clone tracing using single-cell sequencing. Nature 556, 108–112.

Amat, F., Lemon, W., Mossing, D. P., McDole, K., Wan, Y., Branson, K., Myers, E. W. and Keller, P. J. (2014). Fast, accurate reconstruction of cell lineages from large-scale fluorescence microscopy data. Nat. Methods 11, 951–958.

Bouldin, C. M., Snelson, C. D., Farr, G. H. and Kimelman, D. (2014). Restricted expression of cdc25a in the tailbud is essential for formation of the zebrafish posterior body. Genes Dev. 28, 384–395.

Choi, H. M. T., Beck, V. A. and Pierce, N. A. (2014). Next-Generation *in Situ* Hybridization Chain Reaction: Higher Gain, Lower Cost, Greater Durability. ACS Nano 8, 4284–4294.

Duboule, D. (1994). Temporal colinearity and the phylotypic progression: a basis for the stability of a vertebrate Bauplan and the evolution of morphologies through heterochrony. Development 1994,.

Gont, L. K., Steinbeisser, H., Blumberg, B. and de Robertis, E. M. (1993). Tail formation as a continuation of gastrulation: the multiple cell populations of the Xenopus tailbud derive from the late blastopore lip. Development 119, 991–1004.

Handrigan, G.R. (2003). Concordia discors: duality in the origin of the vertebrate tail. J. Anat. 202, 255–267

Henrique, D., Abranches, E., Verrier, L. and Storey, K. G. (2015). Neuromesodermal progenitors and the making of the spinal cord. Development 142, 2864–2875.

Holmdahl DE (1925a) Die erste Entwicklung des Körpers bei den Vögeln und Säugetieren, inkl. dem Menschen, besonders mit Rücksicht auf die Bildung des Rückenmarks, des Zöloms und der entodermalen Kloake nebst einem Exkurs über die Entstehung der Spina bifida in der Lumbosakral region I. Gegenbaurs Morph. Jahrb. 54, 333–384.

Holmdahl DE (1925b) Die erste Entwicklung des Körpers bei den Vögeln und Säugetieren, inkl. dem Menschen, besonders mit Rücksicht auf die Bildung des Rückenmarks, des Zöloms und der entodermalen Kloake nebst einem Exkurs über die Entstehung der Spina bifida in der Lumbosakral region II-V. Gegenbaurs Morph. Jahrb. 55, 112–208.

Holmdahl DE (1925c) Experimentelle Untersuchungen über die Lage der Grenze zwischen primärer and © Anatomical Society of Great Britain and Ireland 2003 sekundärer Körperentwicklung beim Huhn. Anat. Anz. 59, 393–396.

Junker, J. P., Spanjaard, B., Peterson-Maduro, J., Alemany, A., Hu, B., Florescu, M. and van Oudenaarden, A. (2016). Massively parallel whole-organism lineage tracing using CRISPR/Cas9 induced genetic scars. bioRxiv 56499.

Kanki, J. P. and Ho R. K. (1997). The development of the posterior body in zebrafish. Development 124, 881–93.

Martin, B. L. and Kimelman D. (2009). Wnt Signaling and the Evolution of Embryonic Posterior Development. Curr. Biol. 19, R215–R219.

Martin, B. L. and Kimelman D. (2012). Canonical Wnt signaling dynamically controls multiple stem cell fate decisions during vertebrate body formation. Dev. Cell 22, 223–32.

Niehrs, C. K. and C. (2001). AP neural patterning by a Wnt gradient. 1–13.

Nieuwkoop, P. D. and Nigtevecht, G. V Neural Activation and Transformation in Explants of Competent Ectoderm under the Influence of Fragments of Anterior Notochord in Urodeles.

Row, R. H., Tsotras, S. R., Goto, H. and Martin, B. L. (2016). STEM CELLS AND REGENERATION The zebrafish tailbud contains two independent populations of midline progenitor cells that maintain long-term germ layer plasticity and differentiate in response to local signaling cues. 244–254.

Shah, G., Thierbach, K., Schmid, B., Reade, A., Roeder, I., Scherf, N. and Huisken, J. (2017). Pan-embryo cell dynamics of germlayer formation in zebrafish. bioRxiv 173583.

Steventon, B. and Martinez Arias A. (2017). Evo-engineering and the cellular and molecular origins of the vertebrate spinal cord. Dev. Biol. 432,.

Steventon, B., Duarte, F., Lagadec, R., Mazan, S., Nicolas, J.-F. and Hirsinger, E. (2016). Species tailoured contribution of volumetric growth and tissue convergence to posterior body elongation in vertebrates. Development 1732–41.

Tucker AS, Slack JMW (1995a) Tailbud determination in the vertebrate embryo. Curr. Biol. 5, 807–813.

Tucker AS, Slack JMW (1995b) The Xenopus tail-forming region. Development 121, 249–262.

Tzouanacou, E., Wegener, A., Wymeersch, F. J., Wilson, V., Nicolas, J.-F., Beddington, R. S., Rashbass, P., Wilson, V., Bellomo, D., Lander, A., et al. (2009). Redefining the progression of lineage segregations during mammalian embryogenesis by clonal analysis. Dev. Cell 17, 365–76.

Wolff, C., Tinevez, J.-Y., Pietzsch, T., Stamataki, E., Harich, B., Preibisch, S., Shorte, S., Keller, P. J., Tomancak, P. and Pavlopoulos, A. (2017). Reconstruction of cell lineages and behaviors underlying arthropod limb outgrowth with multi-view light-sheet imaging and tracking. bioRxiv 112623.

Vogt W (1926) Ueber Wachstum und Gestaltungsbewegungen am hinteren Körperende der Amphibien. Anat. Anz. 61, 62–75.

Wymeersch, F. J., Huang, Y., Blin, G., Cambray, N., Wilkie, R., Wong, F. C. K., Wilson, V., Al-Kofahi, Y., Lassoued, W., Lee, W., et al. (2016). Position-dependent plasticity of distinct progenitor types in the primitive streak. Elife 5, e10042.

